# Can upwelling regions be potential thermal refugia for marine species during climate warming? The case of Mexico

**DOI:** 10.1101/2023.12.05.570267

**Authors:** Luis Enrique Angeles-Gonzalez, Josymar Torrejón-Magallanes, Luis Osorio-Olvera, Otilio Avendaño, Fernando Díaz, Carlos Rosas

## Abstract

Anthropogenic climate warming will change the thermal habitats of marine species worldwide. Species are expected to migrate to higher latitudes as warming intensifies since behavioral and physiological mechanisms have been adapted to maximize fitness under a specific range of temperatures. However, given the possible intensification of upwelling ecosystems, they may act as potential thermal refugia under climate warming, which could protect the diversity of the marine ecosystems, making them important regions for marine resource management. This research aimed to predict the effects of climate warming on commercial and non-commercial marine species (vertebrate and invertebrates) reported in official Mexican documents (> 300 species) based on a thermal niche characterization with the objective to observe if the upwelling regions can act as potential thermal refugia. For this, we considered Representative Concentration Pathway (6.0 and 8.5) scenarios for the present and the future (2040-2050 and 2090-2100). Current and future patterns of suitability, species distribution, richness, and turnover were calculated via ecological niche models using the minimum volume ellipsoids as an algorithm. The results in this study highlight that beyond migration to higher latitudes, some upwelling regions (not all) could be potentially used as some type of “oasis” by marine species refuging from environmental pressure or zones of higher stability of species. Specifically, the upwelling systems in western Baja California and north of Yucatan may be essential regions for future management in Mexico. Nevertheless, it is important to note that climate change acts on numerous ecosystem features, such as trophic relationships, phenology, and other environmental variables not considered here. Future research could test our hypothesis under more realistic simulations. However, migration toward upwelling regions seems to be the most logical selection from all the available ecosystems as climate refugia for marine species beyond movement to higher latitudes or depths.

## Introduction

The anthropogenic climate change (CC) effect on marine ecosystems has been thoroughly studied to understand and predict its potential impacts [1]. The literature suggests that CC could provoke synergistic environmental effects, usually impacting marine organisms negatively [1–3]. Regardless of the synergetic outcome caused by the different environmental stressors, more substantial evidence and the main focus of the literature have been related to the impact of thermal stress on marine organisms [4–9].

This impact is unsurprising since temperature effects can be observed from molecular to population levels, which are reflected at a biogeographical scale [10–12]. Moreover, a sizeable theoretical framework has been developed around temperature and how physiological mechanisms co-define the animals’ fundamental and realized thermal niches [13]. The available quantity of temperature data, either with satellites, buoys or estimated by hydrodynamic models has allowed addressing the biogeographical impacts of climate warming in the thermal habitat worldwide. For instance, organisms’ distribution shifts are consistent with ocean surface temperature changes [9]. Further evidence has shown that alterations in the composition and abundance of global fishery catches could be associated with temperature changes [5,14].

Species distribution has been modeled under CC to predict the potential redistribution changes it may cause [15–19]. Usually, most works have highlighted latitudinal changes in distribution or migration to deeper waters caused by a difference in environmental requirements. Thus, fishing and conservation scientists must consider the long-term effects of climatic variations to deal with change and weather-induced variability. Despite the potential impact of CC, few studies have aimed to evaluate the vulnerability of fishery resources in Mexico. Mexican researchers assessed the potential effect that CC would have on the coastal ecosystems in Mexico’s Atlantic and Pacific regions [20]. These authors concluded that the impact would harm fisheries that contribute 70% of the fishing volume and economic value due to ecosystem alterations. More recently, a bioclimatic model for 128 marine species predicted substantial decreases in catches in the Campeche Bank in Mexico [18].

Similar to the studies above, collaborators modelled potential changes in thermal habitat based on thermal physiological data finding that besides the migration to higher latitudes well reported by the literature, the upwelling regions of the Campeche Bank and Western Baja California in Mexico might act as thermal or climate refugia [21]. Even though latitudinal shift reports may be the dominant observation, literature relates to the CC effect and the role upwelling regions could play as a climatic refugium [15,22–26]. In the context of this study, climate refugia are regions where the majority of species stay or retreat to endure and possibly expand in the face of changing climatic conditions [25,27,28] – in this case, endure temperature increases. Furthermore, refugia might provide crucial information for better marine resource management [28,29].

Coastal upwelling transports cold water from deep ocean waters to the surface. The upwelling intensity and length dictate if the regional environmental conditions vary temporarily or permanently from those neighboring regions. In addition, the supply of cold waters by upwellings is not directly related to climate, allowing regional decoupling of global warming to serve as a refuge, shielding against existing and predicted climate changes [25] – a desired characteristic in regions with the potential of being climate refugia [27]. Evidence has demonstrated that upwellings regions worldwide are intensifying as a product of CC related mainly to the land-sea pressure difference that could strengthen wind patterns [30,31]. That process could partially buffer coastal upwelling ecosystems from CC temperature increases [26,30]. Indeed, if upwellings regions can act as potential climate refugia, it would be worthwhile to prioritize the management of these regions for diversity or resource protection.

Based on the literature, this study aims at two main objectives: (1) predict the effects of climate warming due to CC in Mexican marine species based on a thermal niche characterization. Correlative ecological niche models (ENMs) were used to achieve this goal. Under an ENM approach, the approximation of a fundamental niche (ideally physiological preferences) is based on environmental space and locations where species are present. Consequently, the ENM can be projected to the geographic areas of interest to generate a species distribution model (SDM) [32]; (2) observe if upwelling regions act as potential climate refugia under CC, protecting the diversity of the marine ecosystems.

## Material and methods

### Study region

Mexico’s Territorial Sea and Exclusive Economic Zone (EEZ) spans 3,149,920 km^2^, with 2,320,380 km² corresponding to the Pacific Ocean and 829,540 km² to the Gulf of Mexico and the Caribbean Sea [33]. The country’s coastline, excluding the islands, measures 11,122 km, with 7,828 km along the Pacific and Gulf of California, and 3,294 km along the Gulf of Mexico and the Caribbean Sea [34]. In coastal upwelling ecosystems, the wind carries subsurface water to the surface by forcing surface water to move offshore. Sub-superficial waters usually have lower temperatures, creating a colder habitat than the regions around [35,36]. Six significant upwelling regions are present in Mexico: (1) North of the Yucatan Peninsula; (2) Campeche Bay; (3) nearby Gulf of Tehuantepec; (4) Mexican Central Pacific in Cabo Corriente; (5) Gulf of California; and (6) Western Baja California/Baja California Sur (Fig 1) [35].

**Fig 1.**
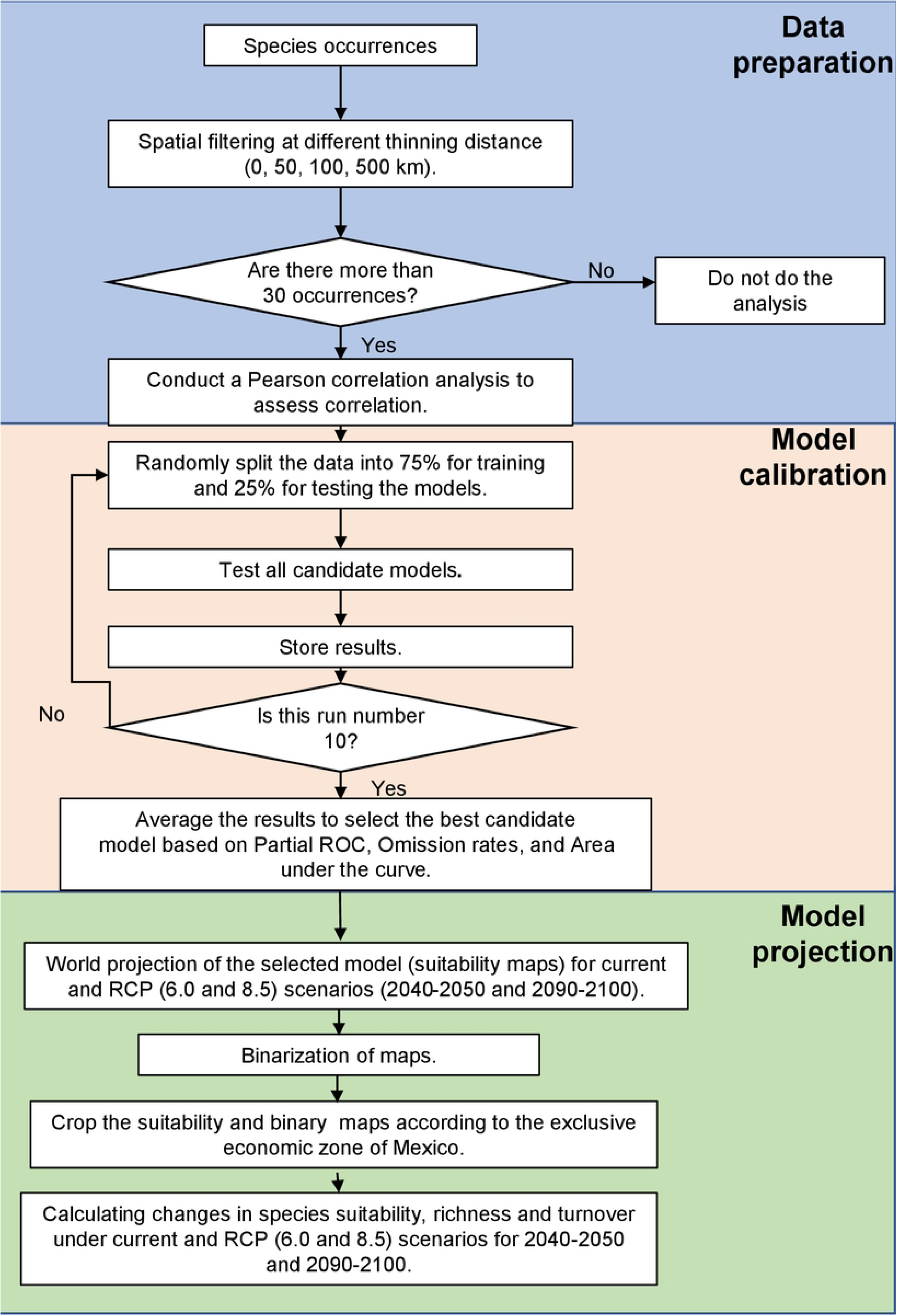
The average daytime sea-surface temperature for the period 2016-2020. The delineated line indicates the Mexican exclusive economic zone. The major upwelling regions are shown as follows: (**1**) North of the Yucatan Peninsula; (**2**) Campeche bay; (**3**) nearby Gulf of Tehuantepec; (**4**) Mexican Central Pacific in Cabo Corriente; (**5**) Gulf of California; and (**6**) Western Baja California/Baja California Sur. Regions 1 and 2 jointly represent the Campeche Bank. Data were obtained from the sensor MODIS-Aqua.

In the Yucatan Peninsula, a seasonal upwelling occurs during spring-summer caused by the friction with the continental slope, creating a topographic upwelling and lowering the temperatures in the region [37]. In the Bay of Campeche, two physical processes that show an effect on the bay productivity are the cyclonic gyre and river discharge. Notably, the cyclonic gyre raises the pycnocline, introducing nutrients in the euphotic zone and creating an upwelling-like zone [36]. In the Gulf of Tehuantepec, winds that occur from May to November derive from the North Winds of the Gulf of Mexico and pull the water to the south creating an upwelling that reduces temperature, increases salinity, and changes the current circulation [35,36]. Little information exists about the mechanism that drives the upwelling in Cabo Corriente, but upwelling has been noticed to intensify during spring [38]. In the Gulf of California, winds come from the northwest during winter and spring, while in summer and autumn, the southeast gets weaker [39]. These wind patterns cause an upwelling in the region to be more intense in the continental part during winter and spring [40]. Finally, in western Baja California, upwellings are generated due to northeastern spring and summer winds, which draw water offshore, further cooling the California Current – an already cold region as it transports cold water from higher latitudes to lower ones [41].

### Environmental data

The current and future maximum, mean, and minimum surface temperatures (SSTmax, SSTmean, SSTmin) were downloaded from Bio-ORACLE (https://www.bio-oracle.org/). The future environmental variables consisted of the Representative Concentration Pathways (RCP) 6.0 and 8.5 for 2040-2050 and 2090-2100 [42]. The future scenarios from Bio-ORACLE were obtained averaging the Community Climate System Model 4 (CCSM4); the Hadley Centre Global Environmental Model 2 (HadGEM2-ES) and the Model for Interdisciplinary Research on Climate (MIROC5) [42]. The RCP 6.0 and RCP 8.5 projection scenarios assume the most severe warming from all RCP with a temperature increase of ∼1°C from 2040-2050 and ∼3 °C from 2090-2100. The most extreme scenario was selected since the aim was to identify the most vulnerable regions and potential climate refugia in Mexico. Bathymetry data from MARSPEC [43] was also downloaded. It is important to mention that the same bathymetric layer was used for current and future scenarios. The environmental layers have a resolution of about 9 x 9 km per pixel at the Equator. All the variables previously mentioned were considered when developing the ENMs (see below).

### Species selection

The selection of the species for modeling was based on the official federal documents “Carta Nacional Pesquera” published during 2017 and 2023 – a document that provides precise and up-to-date information on where, when, and how much fishing is allowed for commercial aquatic species fished in Mexico in addition to incidentally captured species (by-catch) which may not have commercial importance [44–46]. Our goal was to study patterns of species suitability and distribution changes in Mexico. We selected both commercially and non-commercially important species, including vertebrates and invertebrates.

During the selection process, any taxonomy names no longer accepted or wrongly spelled were corrected using as a reference the World Register of Marine Species (WoRMS-https://www.marinespecies.org/), which provides “an authoritative and comprehensive list of names of marine organisms”. This is important since species misidentification can produce inaccurate biological and ecological information under an ENM approach, which could under or overestimate species richness [47]. Finally, it could potentially setback the correct development of effective policies for ecosystem protections [48–50], which could be troublesome under the CC change context. In total, we identified 383 species on Mexican official federal documents (S1 database).

### Occurrence data

Species records were downloaded from the repositories: Ocean Biodiversity Information System (OBIS), Global Biodiversity Information Facility (GBIF), and VertNet using the packages “*robis”* [51], “*rgbif”* [52] and “*rvertnet”* [53] from the programming language R [54], respectively. The species records were curated considering standard steps used in the ecological niche modeling literature [55]. These steps included removing duplicated and wrongly georeferenced records (i.e. inland records). Furthermore, occurrences associated with environmental outliers were visually screened; if the values were too atypical from the bulk of the data (temperatures or bathymetry), they were deleted.

Oversampling and sighting biases are also problematic in online repositories, since they alter the fundamental niche calculated for species. To reduce this problem, first, only one occurrence was considered per pixel. Later, a spatial filtering technique was applied, allowing us to further minimize such issues using the R package “*spThin”* [56]. The *thin* function applies a random spatial thinning at a distance specified by the user. No universal accepted distance exists; thus, four databases were created for each species with a different thinning space applied: 0 (i.e., one occurrence by pixel), 50, 100, and 500 km obtaining a total of 1512 databases.

### Algorithm of the correlative ecological niche models

The minimum volume ellipsoid (MVE) algorithm was used as an ENMs tool to represent the species fundamental niche. The election of this algorithm is because empirical and theoretical works assume that fundamental niches have convex shapes, such as ellipsoids [57]; moreover, the distance to the center of an ellipsoid has been shown to be negatively related to fitness attributes, such as population abundance and genetic variation [58–61]. The MVE is a multivariate estimator that builds ellipsoids spanning a predetermined percentage of observations, predicting a centroid and scattering of the data [62]. The centroid and total volume of the ellipsoids are represented by the combination of non-optimal environmental factors (𝒙 = [𝑥_1_,𝑥_2_,…,𝑥_𝑛_]) and optimal centroid (𝝁 = [𝜇_1_,𝜇_2_,…,𝜇_𝑛_]) where the highest suitability occurs. The suitability function of an ellipsoid depends on the Mahalanobis Distance equation:

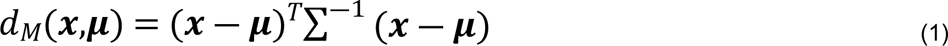

where ∑ is a covariance matrix. Finally, the suitability of *x* is computed as:

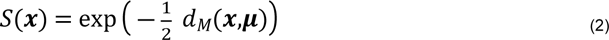

### Calibration process

Before starting the model calibration, we checked for correlations between the environmental variables for each database. Databases with more than 30 occurrences were considered for the entire calibration and projection process. For the correlation analyses, the temperature (SSTmax, SSTmean, and SSTmin) values associated with each occurrence were extracted; posteriorly, a Pearson’s correlation (function *cor* in *base* R) analysis was used to eliminate variables with a value of > 0.7. It is important to emphasize that the variables were deleted according to a previously considered hierarchical order as follows: SSTmax, SSTmean, SSTmin. This hierarchal order was selected because SSTmax values tend to have less dispersed and skewed temperature values than SSTmean and SSTmin. Similarly, the SSTmean were less dispersed and skewed than SSTmin (exploratory figures are stored in figshare - https://figshare.com/s/93dc128799e519b08065). Indeed, given the simplicity of the algorithm MVEs, it is sensible to avoid deviations from symmetry; thus, the hierarchical approach was used trying to minimize such drawbacks [63]. We considered different combinations between surface environmental variables with bathymetry given that only environmental surface models overextended species potential distribution.

Later, each environmental layer remaining after the Pearson correlation analysis was cropped according to a calibration region. The calibration region consisted of a buffer of 500 km around species occurrence for each database. This calibration region is meant to represent an accessibility hypothesis, i.e., areas explored by the species, whether they have been able to settle or not [64]. We extract 10,000 environmental random background points from the calibration region. For the calibration process, the function *ellipsoid_selection* from the “*ntbox*” R package [65] was used under the stratified selection criteria with a repeated subsampling (10 times), where 75% of the occurrences were used to train the MVEs, while the remaining 25% was employed as test data based on all possible environmental variable combinations with the function. For the MVE 97.5% of the data was considered. This inclusion percentage is not affected by sample sizes, improves MVEs when erroneous occurrences registered are about ≤ 20%, and does not perform worse at any level of bias [66]. First, any model without statistical significance (*p* > 0.05) was deleted according to a partial receiver operating characteristic curve (pROC). From the remaining models, we selected the model with the lowest omission rates [67]. Finally, assuming that two or more models had the same performance, the one with the highest area under the curve (AUC) according to the ROC analyses was chosen.

### Suitability maps, richness, and species turnover

The selected models for each species were projected worldwide with the function *ellipsoidfit* from the “*ntbox”* R package. Later, suitability maps were cropped to the Mexican Exclusive Economic Zone (EEZ) for the present (2000-2014) and future scenarios (RCP 6.0 and 8.5, period 2040-2050 and 2090-2100, respectively). Values of 1 would indicate maximum performance and 0 unviable environments. To observe potential changes in the region’s fitness, an overall suitability map was obtained by averaging the suitability maps of all species. This map serves as a proxy for fitness considering that ellipsoids may be a representation of the species convex niche [57,68]. Later, each species suitability map was binarized, i.e., transformed into presence-absence maps, selecting a suitability value that left out the 2.5% of training presences with more atypical environmental combinations.

The taxonomic richness and its changes under CC scenarios were calculated by ‘stacking’, consisting of a sum of binary data frames; i.e., Presence-Absence databases as pixels with values of “1” represent presence and “0” absence. Finally, the species turnover was calculated for all scenarios using the Jaccard distance index in base R (function dist), which represents the proportion of the total species pool shared by the two communities given by the following formula:

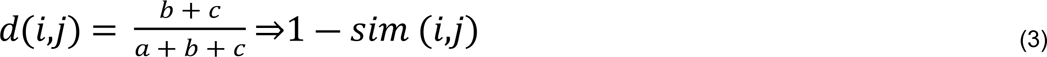

Where *a* is the species shared by pixel in time *i* (present scenario) and *j* (future scenario) represented as 1; *b* is the species present in pixel *i* (1) but not *j* (0); *c* is the species present in pixel *j* but not *i*. These differences were spatially projected to locate regions with higher species turnover (Fig 2). All data and scripts necessary for models replication are added in https://figshare.com/s/93dc128799e519b08065.

**Fig 2.**
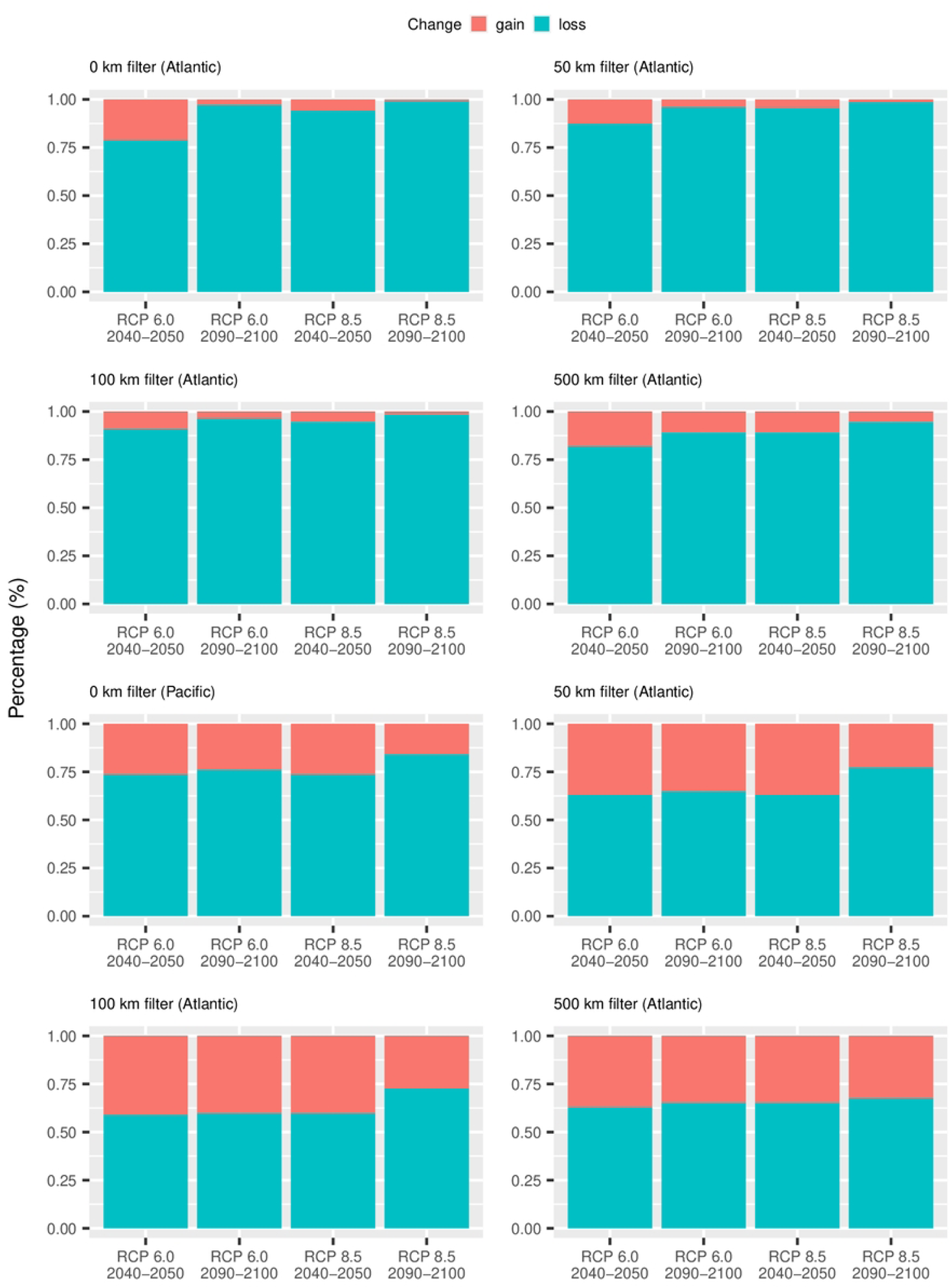
Calibration and projection flow chart. The flow chart shows the entire procedure for obtaining correlative niche models, binarized maps, and calculating richness and turnover (Jaccard distance).

## Results

For most species the selected ENMs have acceptable levels of omission rates (∼0.05) and AUC (>0.9). However, there is an exception with the AUC for the species in the 500 km filter database, where the values tend to be more dispersed reaching values as low as 0.5 (S1 Fig 1). Based on suitability values, we found that all warming scenarios have a negative impact on most marine species in Mexico’s EEZ. This effect is more prominent in the Atlantic region, where the RCP 8.5 scenario (2090-2100) can lead to a decline in the suitability of over 90% of the species. In comparison, the Pacific region may see a decrease in suitability for 80% of the species under the same scenario (Fig 3) with values fluctuating between 60% to 80% for other scenarios.

**Fig 3.**
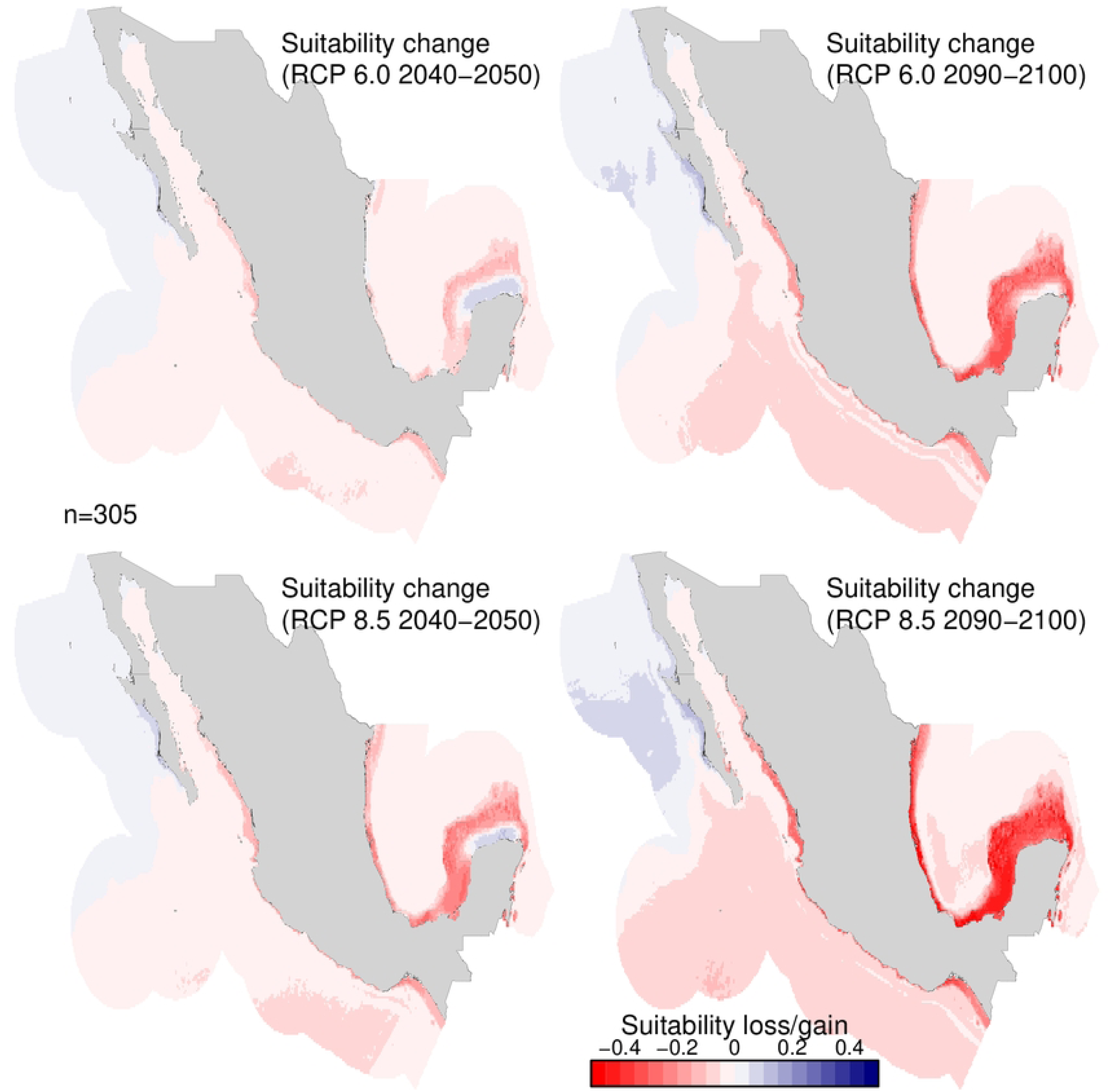
Proportion of species that gain or lose suitability in the Atlantic and the Pacific region under different scales of data filtering, 0 km (occurrence by pixel), 50 km, 100 km, and 500 km.

Indeed, as warming scenarios intensify, lower suitability values are reported (S1 Fig 1.). This is more noticeable in scenarios experiencing intense higher warming, such as in the RCP 6.0 and RCP 8.5 scenarios from 2090-2100. The substraction between average suitability map shows that not all upwelling regions act as thermal refugia. During all warming scenarios and all filtered data (0 km, 50 km, 100 km, and 500 km), there was a reduction in suitability for most species in the Campeche Bay, the Caribbean, and Western Mexico in the Pacific, both in the Gulf of Tehuantepec and the Gulf of Baja California. Conversely, a slight temperature rise (≤ 2 °C) increased the suitability in the north of the Yucatan Peninsula and western Baja California. Further temperature stress reduced species thermal habitat in the Yucatan Peninsula (∼3 °C), with a considerable quantity of species reaching suitability values of almost zero. In contrast, the western and northwestern regions of the Baja California Peninsula still increase suitability (Fig 4 and S1 Fig 1.).

**Fig 4.**
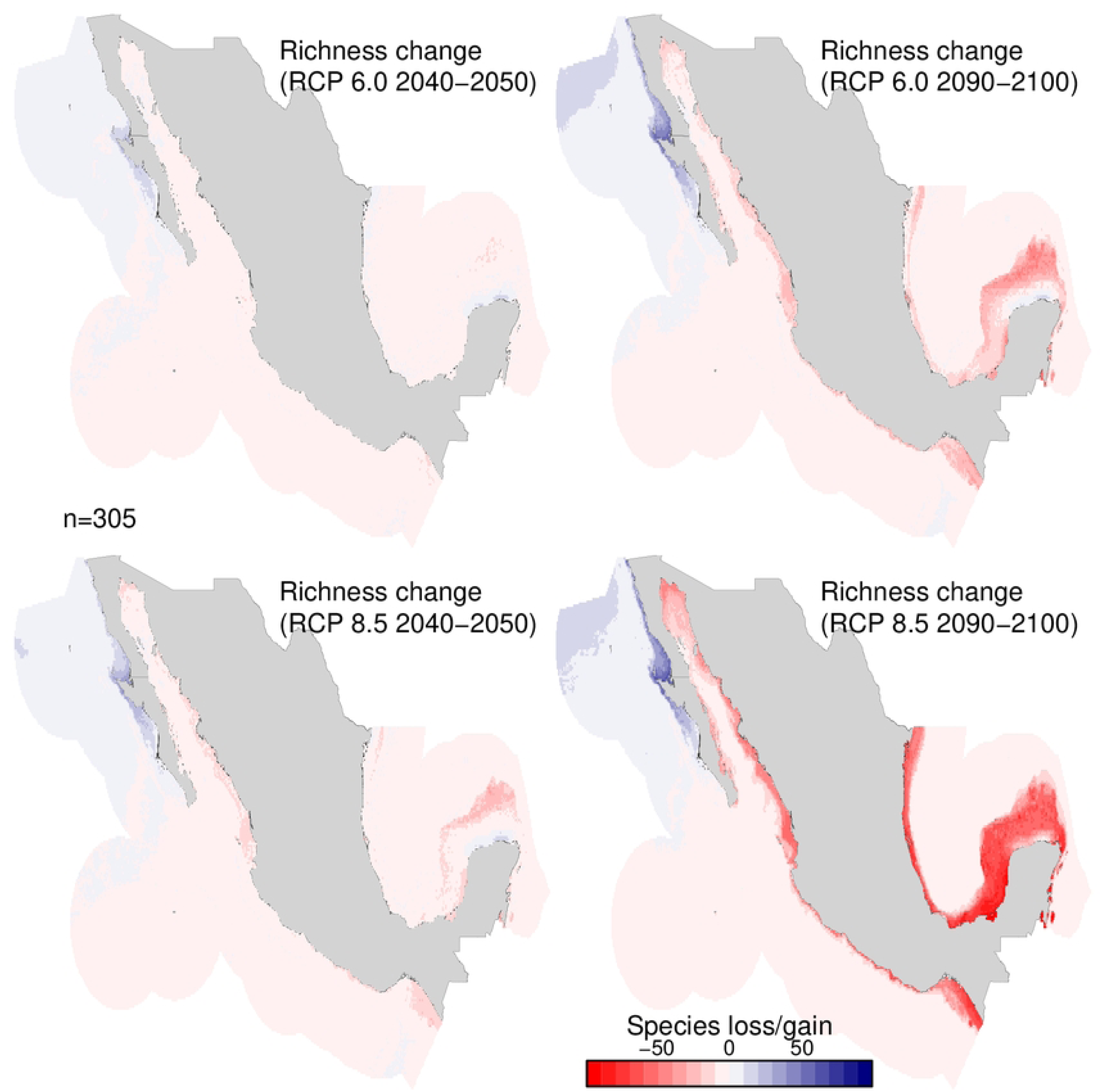
Average suitability of marine species in Mexico’s exclusive economic zone under different climate change scenarios with a spatial filter of 50 km distance.

Patterns of richness change and species turnover follow similar trends as suitability, where temperature rises could momentarily increase richness in the upwelling regions of Yucatan (≤ 2 °C). In addition, according to the Jaccard distance, both Yucatan and south of Baja California act as a buffer area in species composition since the Jaccard distance had values around 0.30 in most scenario (Fig 5 and 6 – S1 Fig.). However, changes in the northwestern region occur in Baja California, likely due to increased richness and the arrival of new species from lower latitudes, even in the RCP 8.5 (2090-2100) scenario. In the other regions, temperature increases (mean temperature ∼4 °C) in the RCP 8.5 (2090-2100) may bring a strong spatial redistribution of species with significant reductions in richness (with regions losing around 50% of species) and a turnover (∼0.5) in most of Mexico. Only for 500 km filter, turnover effects are weak (S1 Fig.).

**Fig 5.**
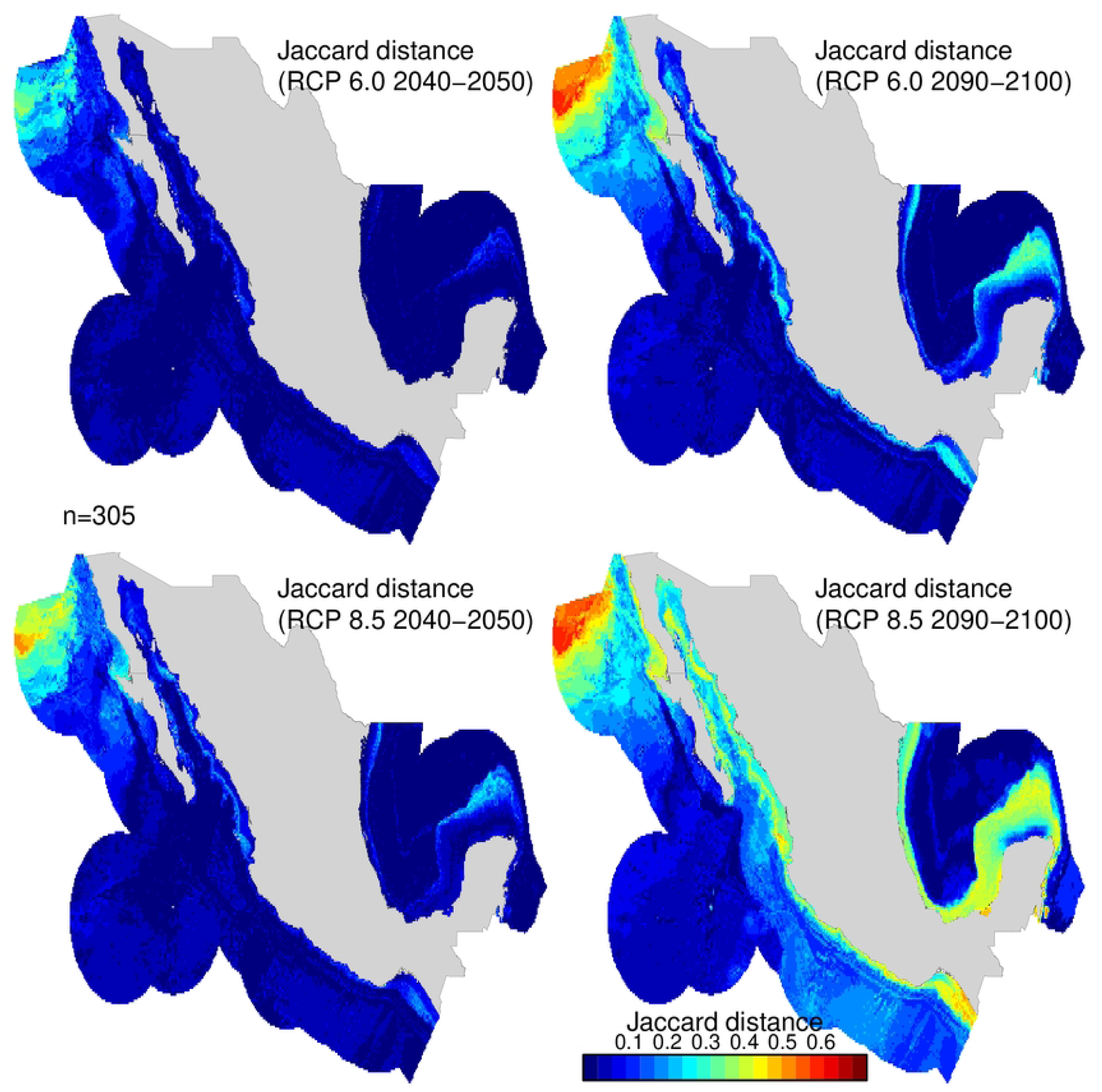
Spatio-temporal loss and gain of marine species in Mexico’s exclusive economic zone under different climate change scenarios with a spatial filter of 50 km distance.

**Fig 6.**
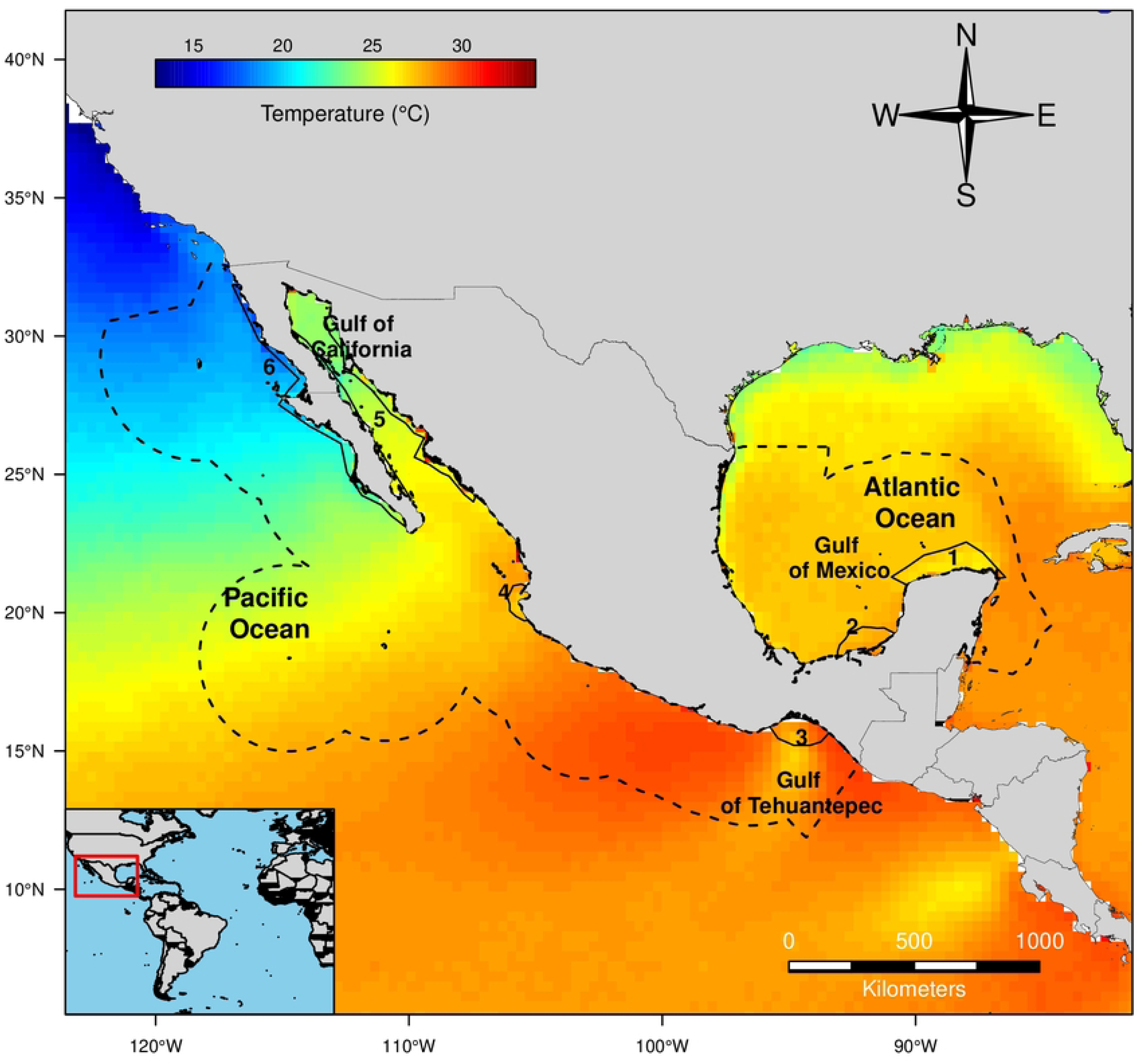
Spatio-temporal turnover of marine species under different climate change scenarios in the exclusive economic zone of Mexico for species thinned spatially at a thinning distance of 50 km.

## Discussion

Studies relating to species vulnerability in Mexico are scarce making our results harder to compare since most studies are usually at a regional scale [15,69–72]. Even so, a few studies have tried to predict the vulnerabilities of fisheries throughout the country [18,20]. This study found spatial patterns of changes in suitability, ecological richness, and turnover through the climate change scenarios; however, their magnitude was related to the warming degree. Even though a direct measurement of abundance was not considered in our work, evidence has shown that estimations via ENMs may reflect important fitness parameters of the population, such as abundance and genetic diversity [59–61,73–75]. In this sense, the results of ENMs can be useful in identifying areas where species abundance may either increase or decrease. Based on this information, we can hypothesize that the regions most vulnerable to temperature increases are the Mexican Caribbean, the western region of Mexico in the Pacific, the Gulf of California, and the west of the Bank of Campeche in the Atlantic.

Our results partially agree with works of our colleagues in the region [18], who noticed that fishery catches could significantly decrease in the Campeche Bank and the Caribbean. Nevertheless, they fail to mention that the upwelling region north of the Yucatan Peninsula could mitigate the decline in catches caused by warming, potentially enhancing species suitability and richness, at least under mild scenarios. More noticeable differences are observed when compared to the Pacific region, as they report potential increases in fishery yields in the central and southern areas. Conversely, in our work, a reduction in species richness, high species turnover, and an overall loss of suitability was usually observed. Nevertheless, the upwelling region nearby the California Current do increases in richness altough it shows a change in species composition according to the Jaccard analysis.

It is noteworthy that only the Yucatan Peninsula and California Current upwelling regions can function as thermal refuge zones in Mexico according to our results. One possible reason for the concentration of species at the north of the Yucatan Peninsula could be the intensity of the upwelling and the protection it provides compared to surrounding waters. After analyzing the environmental layers of Bio-ORACLE, we found that temperatures in the area are on the north of Yucatan average 1 to 3 **°**C lower than surrounding regions, while other upwellings regions are typically only 0.25 to 1 **°**C. This lower temperature on average may be helping species to gather in the area. It is difficult to separate the effect of the upwelling from the latitudinal position of the California Current, but hypothetically, it may still offer protection to organisms migrating to higher latitudes.

It is also remarkably that species turnover was observed across Mexico, with the Yucatan region standing as the sole exception for most filtered databases. In western Baja California (a thermal refugia zone detected in this works), the elevated species turnover is likely linked to the influx of tropical species, leading to a discernible tropicalization effect in tha region. Conversely, in other regions, the turnover is associated with a depletion of species. Only when we applied a spatial filter of 500 km were the effects of turnover less pronounced, likely due to the overrepresentation of species that are more tolerant to temperature changes (S1 Fig.). Nevertheless, with the projected temperature increase under the most extreme scenario (RCP 8.5, 2090-2100), even potential climate refuges, such as upwelling regions, may prove insufficient to fully alleviate the adverse effects of climate change [76], as evidenced in the Yucatan Peninsula.

It has been suggested that certain regions such as the Humboldt and Iberian/Canary eastern boundaries, as well as the central Benguela and California subregions, could provide thermal refuge habitats for their ecosystems. This is due to their cool annual SST footprints that have not contracted over the past two decades in a rapidly warming ocean. However, this is true only for certain seasons [77]. Our colleagues’ observations align with our ENMs.

Marine species’ potential movement to upwelling regions is of general interest to marine biologists and decision-makers since it expands the paradigm of responses to CC. Particularly the discussions of CC and warming by this phenomena have been found mainly in space redistribution to higher latitudes (a pattern also noted in this study) or at greater depths. The above is not strange since spatial redistribution changes in literature are currently well documented [4,5,8,9,16,78,79] but could mask other kinds of potential responses that do not follow those patterns [6,7]. For example, Benguela’s upwelling system has no clear patterns regarding the distribution shift provoked by warming. Fifty percent of the species have changed their distribution to higher latitudes, the rest to the Equator; particularly in South Africa, most species in Namibia moved to shallower waters [8] indicating that the warming region may be now have optimal temperatures for the species. This situation is similar to the response predicted for the Yucatan region, where warming provokes species to migrate to shallower waters. In the Southeastern Pacific, simulations also show that *Scomber japonicus* may conglomerate nearby the Peruvian current, an upwelling system under El Niño and climate change scenarios [19].

Even if the number of studies is comparably lower to descriptions of latitudinal or depth changes, considering the innate characteristics of coastal upwelling, these regions could become essential hotspots for conservation [24,25]. Furthermore, the Eastern Boundary Upwelling Systems have been anticipated to deviate significantly from global change in temperature patterns; indeed, upwelling processes are regarded as a source of bias in global temperature change patterns. However, this information is still debatable because the results vary between sources [31]. For instance, it has been proposed that CC may overwhelm local-scale refugia, particularly during the warmest scenarios [23]. This research study shows that most upwelling regions were “overwhelmed” by warming, even the Yucatan region which provides a degree of protection. Regardless, they seem to be the most logical selection from all the available ecosystems as climate refugia beyond migrations to higher latitudes or depths, at least at the short-term.

One example of the effect of upwelling regions acting as climate refugia is the work related to macroalgae [25]. These authors investigated if upwelling regions could be acting as thermal refugia for the brown alga, *Fucus guiryi,* due to the current effects of climate change while studying their biogeographical dynamics and genetic diversity. They stated that *F. guiryi* had already disappeared from sizable areas of non-upwelling coasts with population numbers far more abundant close to upwelling centers. In addition, genetic analysis of the population revealed distinct genetic groups associated with different upwelling systems. Fishery data can also be used to support the climate refugia-upwelling hypothesis. As suggested based on landing records, mayan octopus (*Octopus maya*) might migrate to the upwelling region from Campeche Bay when in the presence of the high temperatures caused by thermal anomalies [22].

Long-term changes in organisms due to the effect of upwellings have also been described. During the Holocene, rates of vertical accumulation were comparable in two neighboring gulfs, the Gulf of Panama (strong upwelling) and the Gulf of Quiriqui (weak upwelling). Seasonal upwelling in the Gulf of Panama, eventually affected the reef development adversely in the late Holocene. The scenario is presently inverted, with seasonal upwelling in the Gulf of Panama cushioning the thermal stress of climate change. Indeed, the Gulf of Panama reefs produce significantly more carbonate than those in the Gulf of Chiriqui [26].

If behavioral and physiological mechanisms have been adopted to maximize fitness under a range of temperatures (thermal niche), species would be expected to assemble around those thermal habitats that allow them to persist and establish thriving populations [2,12]. This situation is possible because thermal tolerance is linked to the ability to supply oxygen to different tissues, allowing species to maximize fitness-related processes [2,3,13]. As a result, CC will affect fishery yields because populations avert environmental stress, since harmful temperatures can negatively alter species fitness or distribution [80].

If our results are accurate, some upwelling regions in Mexico (and likely worldwide) could provide suitable climate conditions to species under a warming environment, enabling their persistence [25,27,76]. These warming-related safe havens have tremendous ecological and evolutionary relevance since they promote biodiversity conservation [25], allow species to avoid local extinction, and retain genetic variation, which could potentially allow species to expand from those hotspots via adaptation. In addition, identifying climate refuges can provide essential knowledge to improve management by identifying key areas for conservation; thus, some areas like upwelling regions should be prioritized for conservation planning, for both diversity and fisheries [29,81].

### Caveats

It is important to consider that CC is projected to impact not only the thermal habitat of species but also numerous ecosystem features, such as trophic relationships and the phenology of processes related to life histories [8,9], which would alter species distribution. Therefore, it is essential to consider that our estimation does not take into account those critical ecological features. We also have yet to examine CC’s effect on species habitats. For example, pollution, sea-level rise, and many other factors further impact marine organisms in Mexico [20]. Thus, the alterations via CC probably may be more pronounced than what has been predicted in this work.

As stipulated, CC has been expected to decrease oxygen concentrations and increase CO_2_, acidifying the oceans; these processes would potentially reduce the energetic budget of marine species and their thermal niche [1,2,82]. The aforementioned ideas are important because the literature points out that even if the upwelling systems worldwide intensify, deoxygenation and acidification may increase [30,31]. This phenomenon has been impacting biological diversity in the upwelling region of the California Current with deoxygenation and acidification altering the biota distribution [83,84]. Lastly, there is uncertainty in our distribution forecasts due to a lack of certainty in the future impacts of CC on coastal temperatures, notably in upwelling regions [31].

We noticed that performance in terms of AUC varied widely in generalist species (S1 Fig.) usually with worldwide distribution (which are overrepresented in the 500 km filter database). This could be because generalist species can survive in a wide range of environmental conditions, which are not easily defined by the available data and a potential reason for the low turnover reported. In contrast, species with more specific environmental preferences may be easier to model [85]. Finally, we observed a significant spatial bias in the occurrence of certain marine species. Data for these species were mostly reported in higher latitudes, while there were few or no reports in online repositories within the Mexican EEZ. These biases decrease the accuracy of our models [86,87].

## Conclusion

This study highlighted the potential use of upwelling regions as climate refugia via mitigation of the climate warming effect acting like some sort of “oasis” under environmental pressure. Such patterns have been observed in few studies, but there is a lack of studies testing the viability of this hypothesis. In addition, not all upwelling systems may act as solid climate refugia. However, some do have the potential; for example, in this research, the upwelling system in western Baja California and north of Yucatan may be important regions for conservation. Future studies could search for evidence of such potential changes; literature, data recompilation, or fossil records could shed light on this assumption. This information could help us to validate or refute our hypothesis. Indeed, this work only accounted for a thermal niche which must be considered an incomplete reconstruction of the niche. Notwithstanding these concerns, climate refugia are expected to provide the best opportunity for species to survive in future climates. Finally, if the hypothesis is viable, CC studies would benefit from developing more studies in these regions, particularly by conservation biologists and decision-makers.

## Acknowledgments

We are grateful to Diana Fischer for editorial services in English.

**Conceptualization:** Luis Enrique Angeles-Gonzalez, Carlos Rosas; Fernando Díaz, Data Curation: Luis Enrique Angeles-Gonzalez, Otilio Avendaño; **Formal Analyisis**: Luis Enrique Angeles-Gonzalez, Josymar Torrejón-Magallanes, Luis Osorio Olvera; **Software:** Luis Enrique Angeles-Gonzalez, Josymar Torrejón-Magallanes, Luis Osorio Olvera; **Writing – Original Draft Preparation:** Luis Enrique Angeles-Gonzalez, Josymar Torrejón-Magallanes, Luis Osorio-Olvera, Otilio Avendaño, Fernando Díaz, Carlos Rosas; **Writing – Review & Editing:** Luis Enrique Angeles-Gonzalez, Josymar Torrejón-Magallanes, Luis Osorio-Olvera, Otilio Avendaño, Fernando Díaz, Carlos Rosas.

## Supporting information

**S1 database. List of vertebrate and invertebrate marine species of commercial and non-commercial interest reported in Mexico’s “Carta Nacional Pesquera”.** The list shows the scientific and common names of the species found in the “Carta Nacional Pesquera” document from Mexico. It also indicates the specific ocean where they are caught and their ocean distribution regions according to GBIF, OBIS, and Vertnet based on the occurrences available on online repositories. The projections were made considering the data from the repositories.

**S1 Fig. Diagnostic plots, suitability violion plots and maps of average suitability change, richness and turnover of marine species in Mexico’s exclusive economic zone under different climate change scenarios and filtered data.**

